# Dynamic forecasting of Zika epidemics using Google Trends

**DOI:** 10.1101/076521

**Authors:** Yue Teng, Dehua Bi, Guigang Xie, Guigang Xie, Yuan Jin, Baihan Lin, Dan Feng

**Affiliations:** Beijing Institute of Microbiology and Epidemiology, Beijing 100071, China; State Key Laboratory of Pathogen and Biosecurity, Beijing 100071, China; Department of Mechanical and Mechatronics Engineering, University of Waterloo, 200 University Avenue West, Waterloo, Ontario N2L 3G1, Canada; State Key Laboratory for Conservation and Utilization of Subtropical AgroYbioresources, The Key Laboratory of Ministry of Education for Microbial and Plant Genetic Engineering, and College of Life Science and Technology, Guangxi University, 100 Daxue Road, Nanning 530004, China; Beijing Institute of Biotechnology, Beijing 100071, China; Computational Neuroscience Program, Department of Psychology, Physics, and Computer Science and Engineering; Institute for Protein Design, University of Washington, Seattle, WA 98195, USA; Division of Standard Operational Management, Institute of Hospital Management, Chinese PLA General Hospital, Beijing 100853, China

**Keywords:** Zika virus,, Google Trends,, modeling,, forecasting,, surveillance

## Abstract

We developed a dynamic forecasting model for Zika virus (ZIKV), based on real-time online search data from Google Trends (GTs). It was designed to provide Zika virus disease (ZVD) surveillance for Health Departments with early warning, and predictions of numbers of infection cases, which would allow them sufficient time to implement interventions. We used correlation data from ZIKV epidemics and Zika-related online search in GTs between 12 February and 25 August 2016 to construct an autoregressive integrated moving average (ARIMA) model (0, 1, 3) for the dynamic estimation of ZIKV outbreaks. The online search data acted as an external regressor in the forecasting model, and was used with the historical ZVD epidemic data to improve the quality of the predictions of disease outbreaks. Our results showed a strong correlation between Zika-related GTs and the cumulative numbers of reported cases, both confirmed and suspected (both *p<0.001*; Pearson Product-Moment Correlation analysis). The predictive cumulative numbers of confirmed and suspected cases increased steadily to reach 148,510 (95% CI: 126,826-170,195) and 602,721 (95% CI: 582,753-622,689), respectively, in 21 October 2016. Integer-valued autoregression provides a useful base predictive model for ZVD cases. This is enhanced by the incorporation of GTs data, confirming the prognostic utility of search query based surveillance. This accessible and flexible dynamic forecast model could be used in the monitoring of ZVD to provide advanced warning of future ZIKV outbreaks.

## Introduction

Zika virus (ZIKV) is transmitted to people primarily by mosquitoes [1]. Prior to 2015, outbreaks had occurred in Africa, Southeast Asia, and the Pacific Islands [2, 3, 4]. In May 2015, the presence of Zika virus disease (ZVD) was confirmed in Brazil. ZIKV has subsequently reportedly been spreading throughout the Americas, with epidemics occurring in many countries [5, 6]. The World Health Organization declared ZIKV, and its suspected link to birth defects, an international public health emergency in February 2016 [7, 8]. Traditional, healthcare-based and government-implemented, ZVD monitoring is resource intensive and slow. Early warning of infectious disease prevalence, when followed by an urgent response, can reduce the effects of disease outbreaks [9]. Surveillance of online behavior, such as queries in search engines, is a potential web-based disease detection system that can improve monitoring [10]. Google Trends has been shown to have the potential to go beyond early detection and forecast future influenza and Dengue outbreaks [11, 12]. Several studies have used autoregressive integrated moving average (ARIMA) models for the forecasting of influenza prevalence from Google Flu Trends [13, 14]. These models assume their residuals are drawn from a Gaussian distribution, and can be applied to count data by using a logarithmic transformation. The real-time nature of GTs surveillance and the demonstrated strong correlation of GTs with infectious disease mean GTs offers a potential tool for timely epidemic detection and prevention [15]. However, the forecasting capabilities of GTs for ZIKV outbreaks remain unknown. In this study, we examined the ZIKV-related GTs temporally correlated with ZVD epidemics, and developed an improved dynamic forecasting method for ZVD activity in the Americas using an ARIMA model to predict future patterns of ZIKV transmission.

## Results

### Correlations between data on ZIKV outbreaks and ZIKV@related GTs

Our analyses used the data from 12 February to 25 August 2016, covering 7 months of reported ZIKV epidemic data (confirmed and suspected cases) and GTs data (Table S1). Alongside the “stepped” increases in the reported confirmed and suspected cases of ZIKV in this epidemic, there were dramatically increased numbers of Zika-related online searches in Google (Figure 1). From 12 May to 19 May 2016, the ZIKA epidemics entered a period of rapid growth with a large cumulative number of reported confirmed cases (increasing from 8,670 to 40,479; Figure 1A). Meanwhile, the cumulative number of reported suspect cases fell from 298,488 to 269,876 (Figure 1B). Until 25 August 2016, a total of 111,333 confirmed cases and 466,815 suspected cases were reported in the Americas. The GTs data show that the volume of dynamic Zika-related online searches continuously grew from February to August 2016 (Figure 1C). We performed Pearson Product-Moment Correlation analyses to examine temporal correlations between the cumulative numbers of reported cases and the accumulative volumes of Zika-related search queries. The result indicated that the data on Zika-related GTs had statistically significant and positive correlations with the cumulative numbers of both confirmed cases (0.78, *p<0.001*) and suspected cases (0.99, *p<0.001*) of ZIKV.

**Figure 1.**
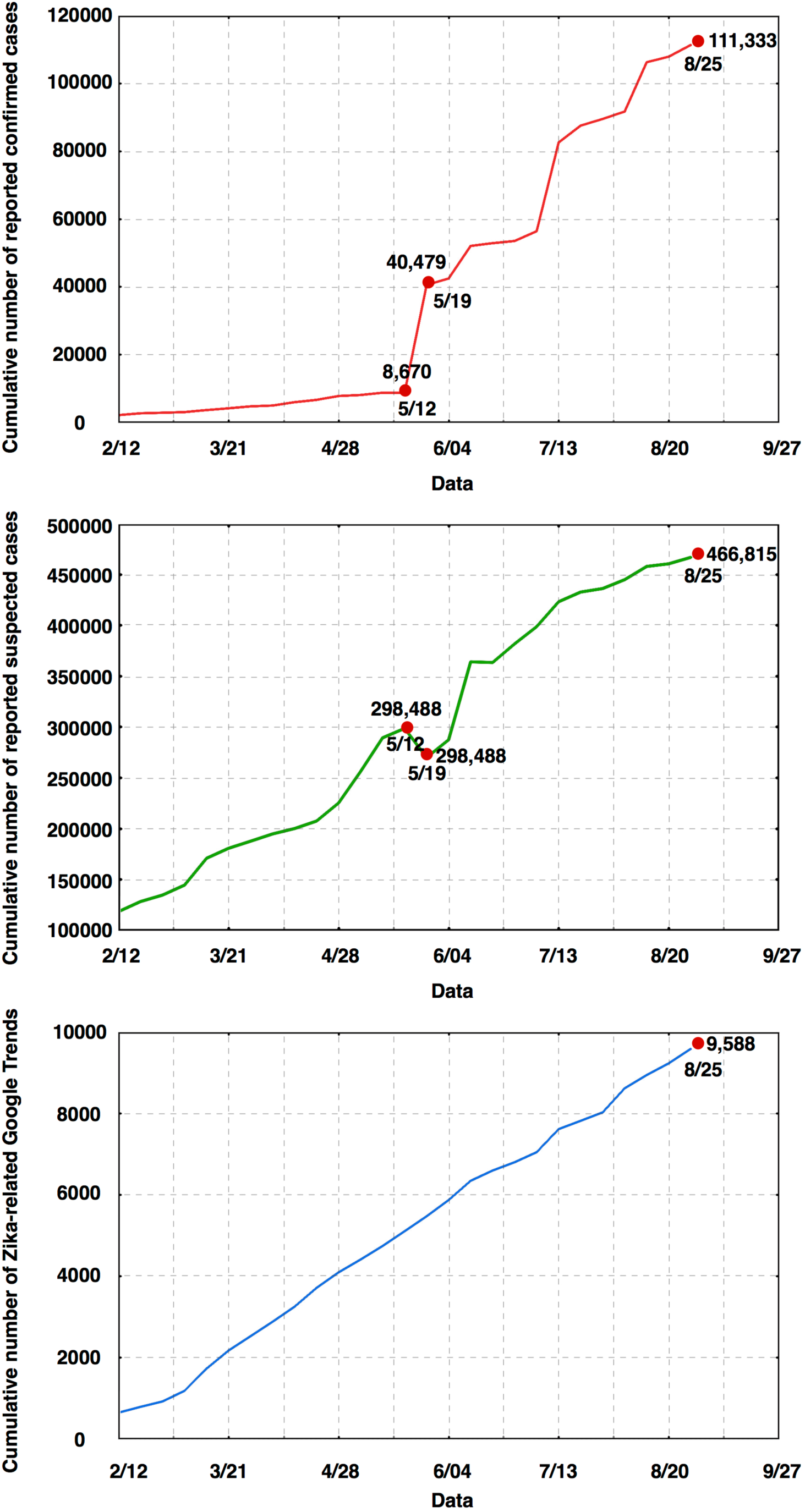
Time series plots of the cumulative number of reported confirmed cases (A), reported suspected cases (B), and Google searches (C) about ZIKV in Americas during ZVD epidemic from 12 February to 25 August 2016.

### Statistical machine learning and reconstructed ARIMA model

Base on the revealed correlation between the data of GTs and the number of reported cases, we split the entire data into training (75%, from 12 February to 28 July 2016) and testing (25%, from 5 August to 25August 2016) sets. We analyzed the training set to observe whether the simulation data is similar to the reported data in testing set using the advanced autoregressive integrated moving average (ARIMA (0, 1, 3)) model. In this reconstructed model, we used the collected online search data as the external regressor in a prediction model to assist the historical ZVD epidemic data in improving the quality of prediction. Using this model with the training data of GTs as a predictor, we found that the cumulative number of reported confirmed case (111,333) in 25 August 2016 in the testing set fell within the 95% confidence interval of the simulated data (93,540 to 127,400) (Figure 2A). However, Figure 2B shows that the number of suspected case (466,815) reported on 25 August 2016 in the testing set was slightly below the lower limit of the 95% confidence interval for the simulated data (476,077 to 525,811). These observations implied that the Google Trends information would improve the prediction of the size of ZIKV outbreaks.

**Figure 2.**
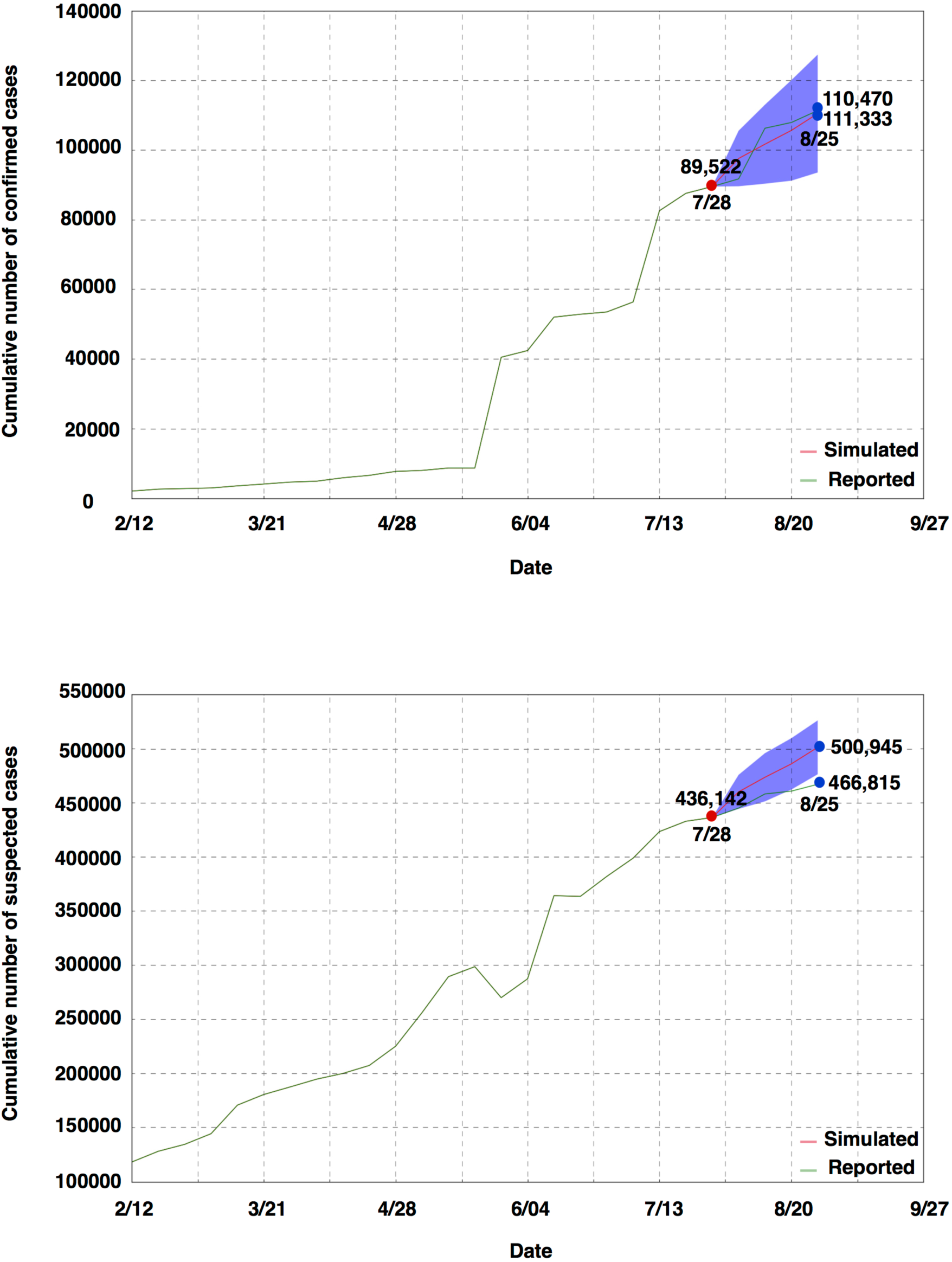
Numbers of reported confirmed cases (A) and suspected cases (B) in the testing set compared with the simulation data by the advanced ARIMA (0, 1, 3) model for training set using the data of Google Trends as the external regressor.

### Dynamic nowcasting of the Google Trends data

As previously described, the Zika-related GTs had a strong correlation with ZIKV associated cases. To improve the prediction of the trend of ZVD epidemics, we estimated the dynamic volumes of Zika-related online searches in Google and used this as a predictor. We passed the available data from GTs to the in-sample baseline ARIMA (0, 1, 3) model to forecast the weekly updated new data from September to October 2016 (Table S2). Figure 3A shows the historical weekly new data, with the estimated weekly new data, and the result predicted that the weekly new data would remain around 350 searches per-week (95% CI: 228-475) for the next 8 weeks until 21 October 2016. Combining the historical and predicted weekly new data of GTs suggested that the forecast cumulative number of GTs would increase to 10996 (95% CI: 10,532-11,459) in 22 September 2016 and might reach 12398 (95% CI: 11,447–13,349) by 21 October 2016 (Figure 3B). These results were used as a forecasting predictor to assist and improve the prediction of ZVD cases in potential future disease outbreaks.

**Figure 3.**
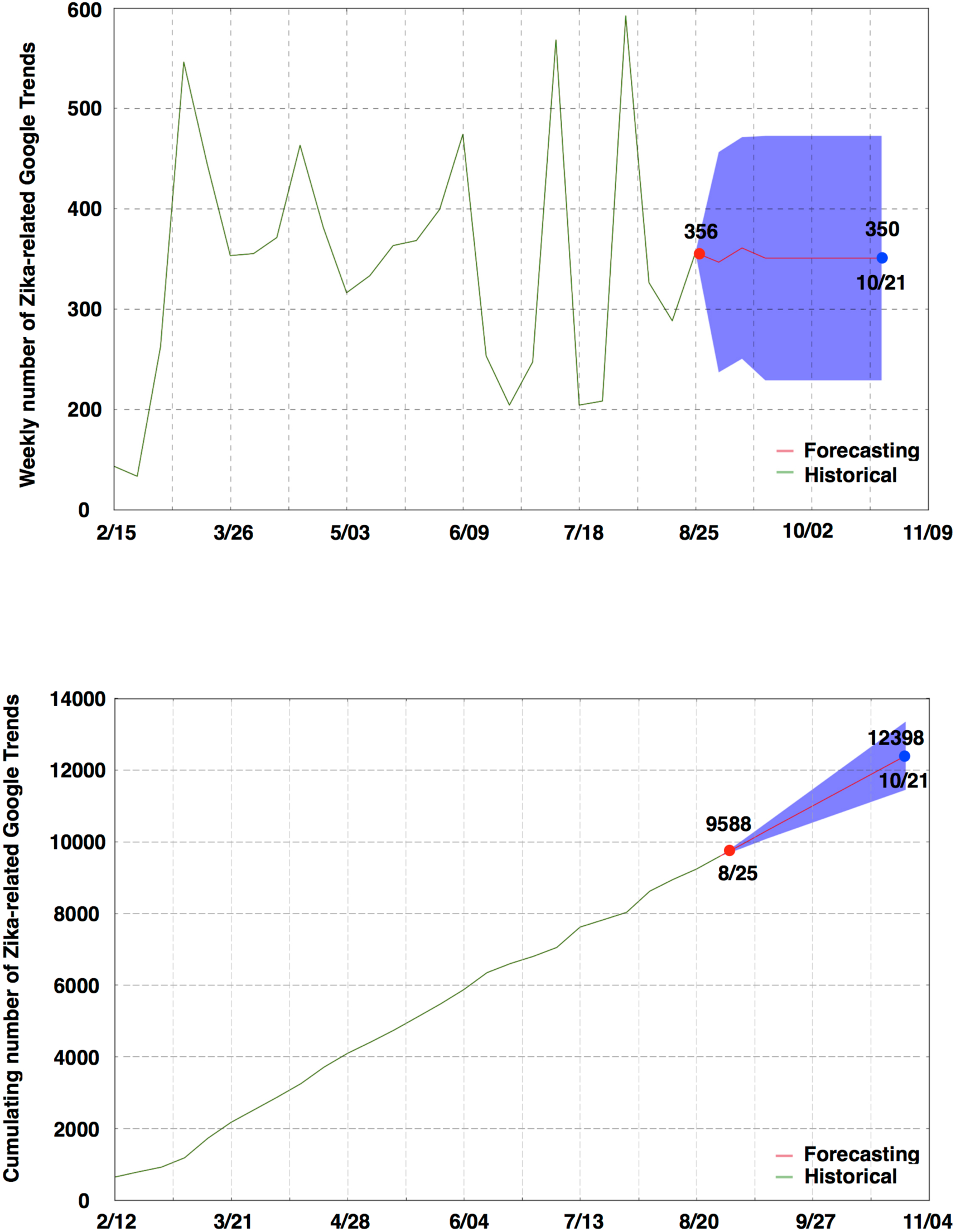
Expected values for cumulative data (A) and weekly updated new data (B) in Google Trends from 2 September to 21 October 2016 (8 weeks) predicted by the in-sample baseline ARIMA (0, 1, 3) model from the Google Trends data covering the first 29 weeks.

### Forecasting the ZIKV outbreaks in Americas during September and October 2016

Based on the data on ZIKV epidemics and Zika-related online searches in GTs between 12 February and 25 August 2016, we used the reconstructed ARIMA (0, 1, 3) model to forecast future ZVD outbreaks in the Americas. In this model, we used the online search data as the external regressor to enhance the forecasting model and assist the historical ZVD epidemic data in improving the quality of the predictions while responding to disease outbreaks. We forecasted the cumulative number of reported ZIKV confirmed cases and suspected cases, respectively, as a simulation of the continuation of the time series. This model with GTs data as the predictor estimated that the forecast cumulative number of confirmed cases would increase to reach 128,826 (95% CI: 113,420-144,231) by 22 September. If current conditions persist, this number would continue to grow to 148510 (95% CI: 126,826-170,195) by 21 October 2016 (Figure 4A). Alongside the constantly increasing number of forecasting-confirmed cases, an increasing number of suspected cases is expected (Figure 4B). The predicted number of suspected cases would grow up to 602,721 (95% CI: 582,753-622,689) on 21 October 2016 (Table S3). These forecasts of ZVD outbreaks suggest that ZIKV disease transmission in the Americas remains intense during September and October 2016.

**Figure 4.**
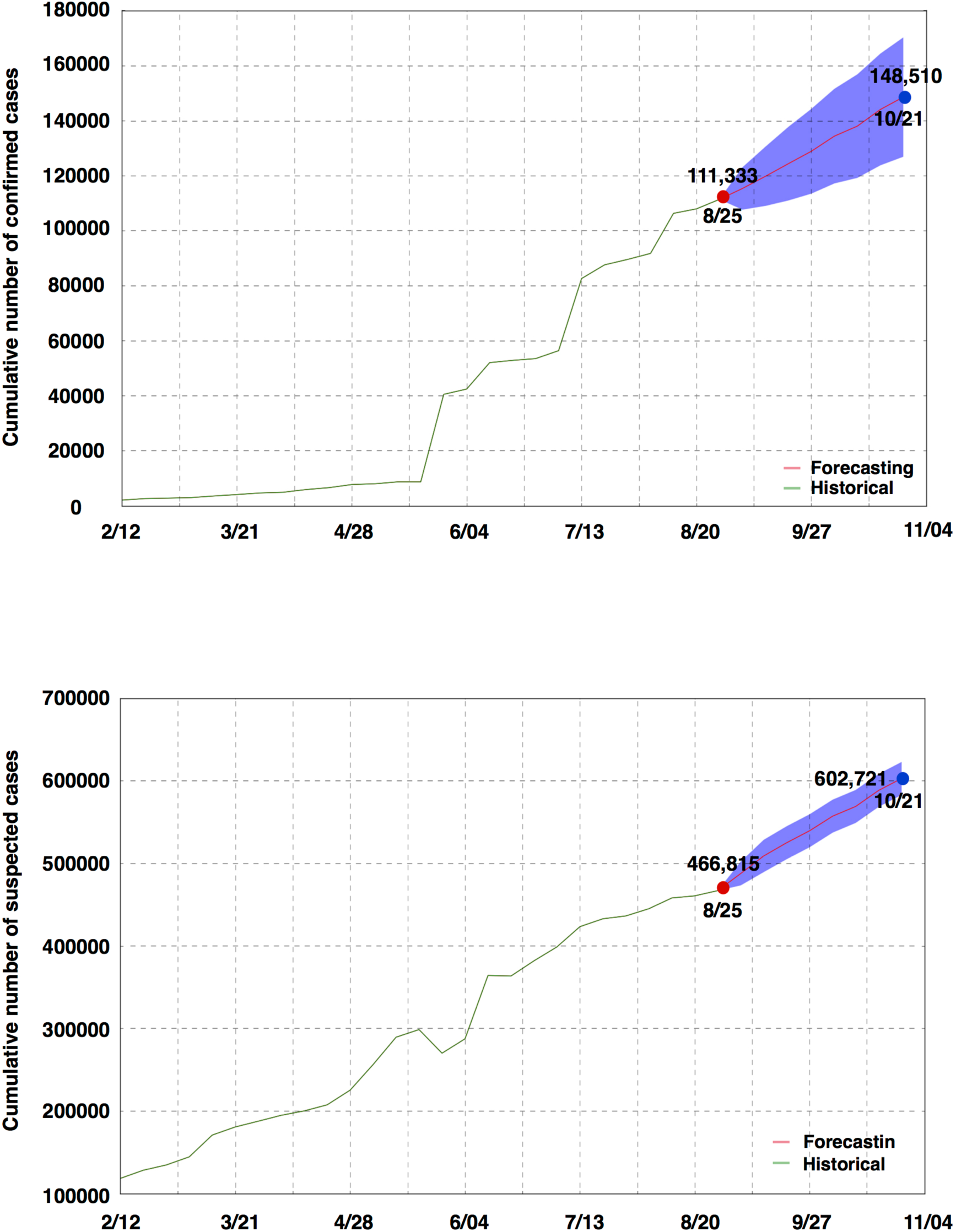
Forecasts of the cumulative number of ZIKV confirmed cases (A) and suspected cases (B) in Americas between 2 September and 21 October 2016 by the advanced ARIMA (0, 1, 3) model, which was improved by aggregating historical logs with estimated data of Zika-related Google Trends as a estimating predictor to estimate ZVD cases.

## Conclusions

ZVD outbreaks are now a common and growing problem worldwide [16, 17, 18]. Delays in traditional surveillance systems limit the ability of public health agencies to respond efficiently to ZIKV epidemics [19]. Because data on GTs are collected and processed in near real-time, online search information produces monitoring data much faster than traditional systems [20, 21]. We first performed correlation analyses to investigate the temporal correlations between data on reported cases of ZVD and ZIKV-related GTs. The result showed that the Zika-related GTs had a strong correlation with both confirmed and suspected cases of ZIKV. Based on the correlation data, the advanced model ARIMA (0, 1, 3) was improved by aggregating historical logs and estimated data of online search queries associated with Zika, as a forecasting predictor, to estimate ZVD cases. The results indicated that the forecasted cumulative number of confirmed and suspected cases would continuously and rapidly grow to reach 148,510 (95% CI: 126,826-170,195) and 602,721 (95% CI: 582,753-622,689), respectively, by 21 October 2016. Although web access is likely to increase in the future, the novel surveillance tool of GTs can also provide dynamic timely information to public health agencies and provide near real-time indicators of the spread of infectious disease [22]. However, to be effective for monitoring disease activity on local geographic areas, it must be considered within the local context of ZIKV transmissibility.

## Materials and Methods

### Data collection and Statistical Analysis

Google Trends, an online tracking system of Internet hit-search volumes (Google Inc.), was used to explore web behavior related to the ZIKV outbreaks. GTs data for ZIKV in Americas was mined from 1 January 2016 to 31 August 2016 to cover the entire period of the ZIKV epidemic, and was downloaded directly from https://www.google.com/trends/explore?date=all&q=zika on 25 August 2016 (data shown in Table S1). The number of ZIKV infected cases in the Americas was retrieved from the PAHO (Pan American Health Organization), available at http://www.paho.org/hq/ (last accessed on 25 August 2016) (Table S1). Numbers of weekly and overall cases of ZIKV were used for the analysis. To quantify the accuracy of GTs relative to reported ZIKV confirmed and suspected cases, we used the Pearson Product-Moment Correlation to assess linear correlation. All calculations were performed in Python 2.7 with the Scipy library.

### Reconstructed ARIMA model

For the time series analysis, we fitted an autoregressive integrated moving average (ARIMA) (0, 1, 3) model by using R version 2.14 (http://www.r-project.org/). The autoregressive integrated moving average (ARIMA) forecasting model in this study was developed from the training and testing sets that were extracted from the data sets. We used the data set including the 7 months from 12 February to 25 August 2016 in the analysis. The data from 12 February to 28 July 2016 were used for training the forecasting model, and the validation was performed on the remaining 4 weeks data. The model was fitted by the standard approach for creating time-series forecasting models. This is described in more detail by Stock & Watson [23]. Using this model selection procedure, we selected one non-seasonal difference term for stationary (d) and three lags of moving average terms (q), resulting in a model of ARIMA (0, 1, 3).

To predict the future values, the developed ARIMA model was fitted to the entire data from 12 February to 25 August 2016 and used to forecast over a time span of 8 weeks, covering September and October 2016. The predictor values were based on predictions of weekly updated GTs data for the predicted periods (8 weeks) from the ARIMA model fitted to the historical data of GTs. We then summed the weekly updated GTs data to obtain estimates of the cumulative online searching data, and this was used as the predictor in this model to forecast the number of cumulative confirmed and suspects ZIKV cases.

## Acknowledgements

We thank Drs. Wuchun Cao and Yigang Tong in State Key Laboratory of Pathogen and Biosecurity for discussions and our colleagues in the Beijing Institute of Microbiology and Epidemiology for help in technical assistance. This work was supported by grants from the State Key Laboratory of Pathogen and Biosecurity Program (No. SKLPBS1408 and No. SKLPBS1451).

## Author Contributions

The manuscript was written by Y.T. and D.H.B.; Data analyses were performed by D.H.B., C.G.X., B.H.L. and Y.T.; The study was designed by Y.T., D.H.B., Y.J., and D.F.

## Author Information

The authors declare no competing financial interests. Correspondence and requests for materials should be addressed to Y.T.

---

**Table S1.** Cumulative number of reported confirmed cases, reported suspected cases, and Google searches about ZIKV in the Americas during the ZVD epidemic of 12 February to 25 August 2016 (29 weeks).

| Date | Number of Confirmed Cases | Number of Suspected Cases | Number of Google Trends |
| --- | --- | --- | --- |
| 2/12/16 | 2048 | 118208 | 637 |
| 2/18/16 | 2601 | 128087 | 780 |
| 2/25/16 | 2765 | 134460 | 913 |
| 3/3/16 | 2943 | 144290 | 1175 |
| 3/10/16 | 3542 | 170683 | 1721 |
| 3/17/16 | 4072 | 180493 | 2166 |
| 3/24/16 | 4629 | 187473 | 2519 |
| 3/31/16 | 4883 | 194618 | 2874 |
| 4/7/16 | 5869 | 199922 | 3245 |
| 4/14/16 | 6560 | 207312 | 3708 |
| 4/21/16 | 7698 | 225088 | 4089 |
| 4/28/16 | 7982 | 256076 | 4405 |
| 5/5/16 | 8672 | 289233 | 4738 |
| 5/12/16 | 8670 | 298488 | 5101 |
| 5/19/16 | 40479 | 269876 | 5469 |
| 5/26/16 | 42399 | 287265 | 5868 |
| 6/2/16 | 52003 | 363990 | 6342 |
| 6/9/16 | 52827 | 363344 | 6595 |
| 6/16/16 | 53473 | 381726 | 6799 |
| 6/24/16 | 56350 | 398626 | 7046 |
| 7/14/16 | 82559 | 423160 | 7614 |
| 7/21/16 | 87542 | 432693 | 7818 |
| 7/28/16 | 89502 | 436142 | 8026 |
| 8/5/16 | 91692 | 444884 | 8618 |
| 8/11/16 | 106252 | 457894 | 8944 |
| 8/18/16 | 107888 | 460496 | 9232 |
| 8/25/16 | 111333 | 466815 | 9588 |

**Table S2.** Expected total and weekly new updated volumes of Zika-related online searches in Google Trends between 2 September and 21 October 2016 (8 weeks) using the ARIMA (0, 1, 3) model.

| Date | Predicted weekly new Google Trends | Expected total Google Trends |
| --- | --- | --- |
| 9/2/16 | 346 with a 95% CI: (237 - 456) | 9934 with a 95% CI: (9825 - 10044) |
| 9/8/16 | 361 with a 95% CI: (250 - 471) | 10295 with a 95% CI: (10075 - 10515) |
| 9/15/16 | 350 with a 95% CI: (228-475) | 10646 with a 95% CI: (10303 - 10988) |
| 9/22/16 | 350 with a 95% CI: (228-475) | 10996 with a 95% CI: (10532-11459) |
| 9/29/16 | 350 with a 95% CI: (228-475) | 11347 with a 95% CI: (10761 - 11932) |
| 10/7/16 | 350 with a 95% CI: (228-475) | 11697 with a 95% CI: (10989 - 12405) |
| 10/14/16 | 350 with a 95% CI: (228-475) | 12048 with a 95% CI: (11218 - 12877) |
| 10/21/16 | 350 with a 95% CI: (228-475) | 12398 with a 95% CI: (11447 - 13349) |

**Table S3.**
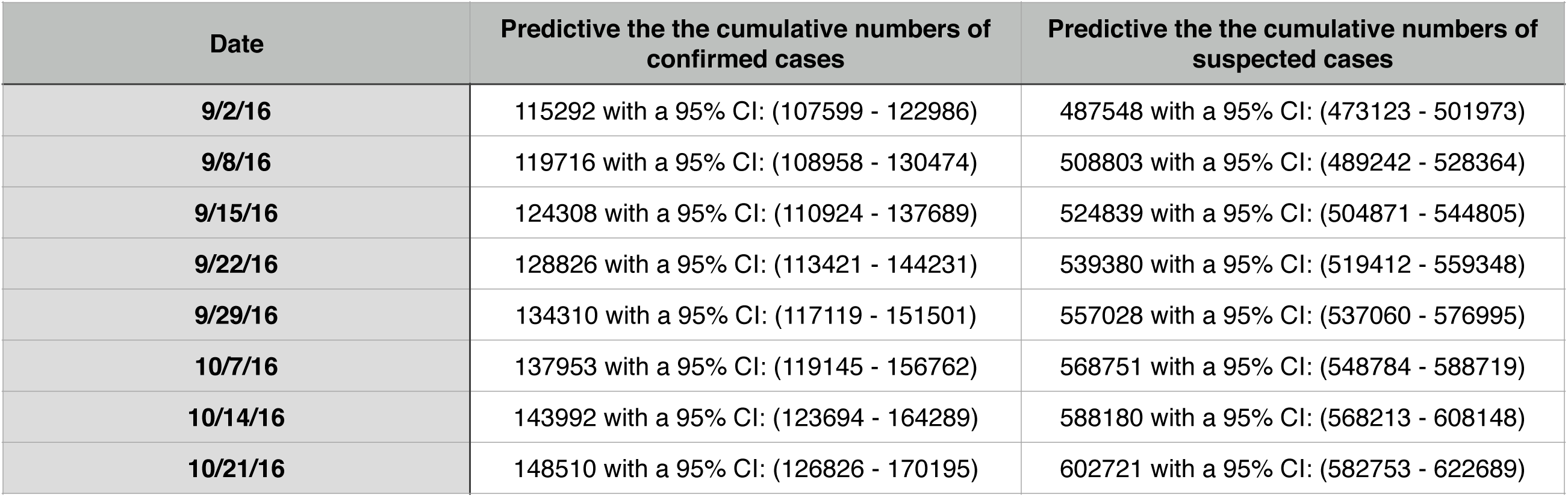
Predictions of the cumulative numbers of confirmed cases and suspected cases from 2 September to 21 October 2016 by the advanced ARIMA (0, 1, 3) model, which was improved by aggregating historical logs with estimated data of Zika-related Google Trends as a estimating predictor to estimate ZVD cases.

| Date | Predictive the the cumulative numbers of confirmed cases | Predictive the the cumulative numbers of suspected cases |
| --- | --- | --- |
| 9/2/16 | 115292 with a 95% CI: (107599 - 122986) | 487548 with a 95% CI: (473123 - 501973) |
| 9/8/16 | 119716 with a 95% CI: (108958 - 130474) | 508803 with a 95% CI: (489242 - 528364) |
| 9/15/16 | 124308 with a 95% CI: (110924 - 137689) | 524839 with a 95% CI: (504871 - 544805) |
| 9/22/16 | 128826 with a 95% CI: (113421 - 144231) | 539380 with a 95% CI: (519412 - 559348) |
| 9/29/16 | 134310 with a 95% CI: (117119 - 151501) | 557028 with a 95% CI: (537060 - 576995) |
| 10/7/16 | 137953 with a 95% CI: (119145 - 156762) | 568751 with a 95% CI: (548784 - 588719) |
| 10/14/16 | 143992 with a 95% CI: (123694 - 164289) | 588180 with a 95% CI: (568213 - 608148) |
| 10/21/16 | 148510 with a 95% CI: (126826 - 170195) | 602721 with a 95% CI: (582753 - 622689) |

